# The iPSC proteomic compendium

**DOI:** 10.1101/469916

**Authors:** Alejandro Brenes, Dalila Bensaddek, Jens Hukelmann, Vackar Afzal, Angus I. Lamond

## Abstract

Induced pluripotent stem cell (iPSC) technology holds great potential for therapeutic and research purposes. The Human Induced Pluripotent Stem Cell Initiative (HipSci) was established to generate a panel of high-quality iPSCs, from healthy and disease cohorts, with accompanying multi-omics and phenotypic data. Here, we present a proteomic analysis of 217 HipSci iPSC lines obtained from 163 donors.

This dataset provides a comprehensive proteomic map of iPSCs, identifying >16,000 protein groups. We analyse how the expression profiles of proteins involved in cell cycle, metabolism and DNA repair contribute to key features of iPSC biology and we identify potential new regulators of the primed pluripotent state. To facilitate access, all these data have been integrated into the Encyclopedia of Proteome Dynamics (www.peptracker.com/epd), where it can be browsed interactively. Additionally, we generated an iPSC specific spectral library for DIA which we deposited in PRIDE along with the raw and processed mass-spectrometry data.

## Main text

A decade ago, Yamanaka and colleagues reported methods allowing the induction of Pluripotent Stem Cells (iPSCs) from human fibroblast cultures^1^. Their report showed that by exogenously expressing a small set of key transcription factors, a somatic cell could be reprogrammed back into a pluripotent state. These reprogrammed cells were shown to have the key features of their physiological embryonic stem cell (ESC) counterparts. Furthermore, since these reprogrammed stem cells can subsequently be differentiated into different somatic cell types, there is currently great interest in using iPSC technology to model human disease mechanisms and study development.

The Human Induced Pluripotent Stem Cells Initiative (HipSci) was established to generate and characterise a large, high-quality panel of human iPSCs, accompanied with detailed multi-omics and phenotypic characterisation. This involved the systematic derivation of iPSCs from many hundreds of healthy volunteer donors, along with disease cohorts, using a standardized and well-defined experimental pipeline that was previously described^2^.

Here, we provide a comprehensive, high-resolution proteomic map of human iPSCs, using lines generated by the HipSci consortium. Data are integrated from the quantitative, mass spectrometry-based analysis of 217 different iPSC lines derived from normal and disease cohorts. We provide a comprehensive proteomic overview of the human iPS cell type and explore how protein expression profiles account for key aspects of iPS cell biology affecting metabolism, cell cycle and DNA repair. Additionally, by comparing protein expression between iPSC lines with either a ‘High’ or ‘Low’ Pluritest^3^ score, we identify potential new regulators of the primed pluripotent cell state.

To provide open access to all these data, the raw and the processed MS files, together with a spectral library for DIA analysis of human iPSCs, have been deposited to the ProteomeXchange^4^ PRIDE^5^ repository (PXD010557), while the processed protein level data have been integrated into the Encyclopedia of Proteome Dynamics^6^, where it can be explored and analysed interactively.

## Results

### Comprehensive Proteomic map of human iPSCs

To provide a comprehensive overview of the human iPSC proteome, we used mass spectrometry-based (MS) quantitative proteomics to analyse 217 separate cell lines derived from 163 different donors. The iPSCs were all derived from human skin fibroblasts and reprogrammed as described previously^2^ (illustrated in Fig. 1a).

**Figure 1.**
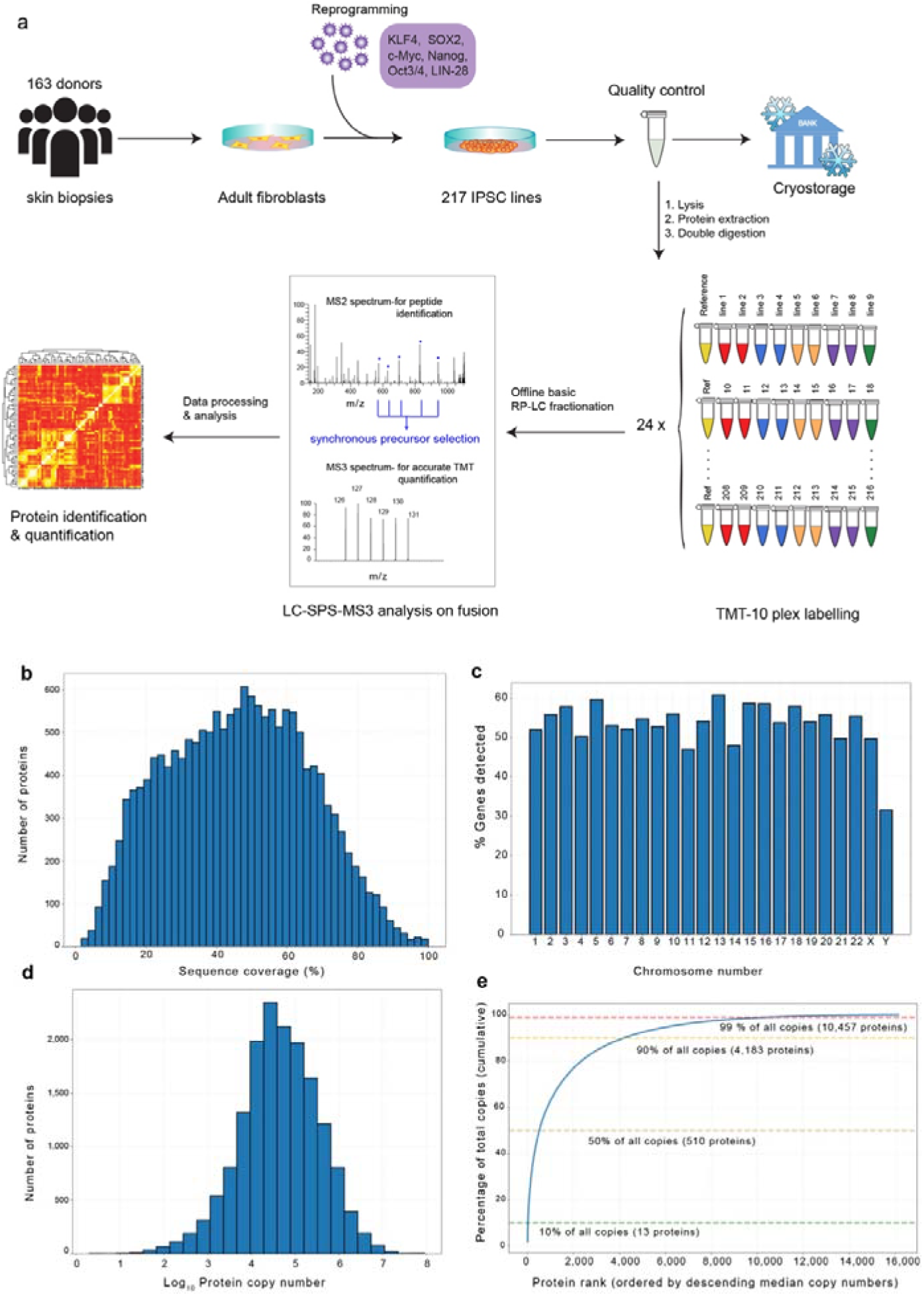
iPSC Proteome: **(a)** The HipSci proteomics workflow from reprogramming to identification and quantification. **(b)** Histogram showing the sequence coverage for all proteins quantified. **(c)** Percentage of protein coding genes detected per chromosome. **(d)** Histogram showing the distribution of the median protein copy number across all lines. **(e)** Cumulative abundance plot showing the percentage of total copy numbers accumulated by protein rank, ordered by descending copy numbers

To maximise throughput for the MS analysis, we combined high mass-accuracy MS with tandem mass tagging (TMT). This allowed the multiplexing of up to 10 cell lines in a single batch. The study was designed so that each batch contained 9 different iPSC lines and one common reference cell line, as shown in Fig.1a. The proteomic analysis of the 217 human iPSC lines was thus divided into 24 batches of 10plex-TMT experiments.

To maximize proteome coverage, a two-dimensional LC-MS strategy was used (see Methods). The first dimension was an off-line, basic reverse-phase HPLC fractionation step, separating the digested and TMT-labelled extracts from each iPSC line into 24 fractions. Each fraction was then analysed using online LC-MS on an Orbitrap Fusion MS. The resulting MS data were subsequently processed and quantified using the MaxQuant^7^ suite.

In total, >16,000 protein groups were detected at 5% FDR (PSM and protein level). This corresponds to detecting the expression of proteins encoded by over 10,500 different genes, representing 52% of all the human protein coding genes annotated within SwissProt. The depth of analysis achieved corresponds to a median protein sequence coverage of ∼46% across all proteins (Fig.1b). Furthermore, this depth of protein coverage was relatively constant across most chromosomes. For the majority of chromosomes, we detected >50% of their protein coding genes expressed in iPSCs (Fig. 1c).

It is known from previous studies on differentiated human cells and cancer cell lines that there is a wide dynamic range of protein expression levels^8^. Understanding the copy numbers of specific proteins expressed in a cell can provide important insights into the metabolism and physiological state of each cell type. To investigate protein expression levels in the iPS cell lines, we estimated protein copy numbers using the ‘proteomic ruler’ approach^9^, which calibrates protein expression relative to the level of histone proteins detected. This is well suited to analysis of the HipSci iPS lines, which are known to have near identical DNA content with little or no copy number variation^2^.

Using the proteomic ruler, we detected wide differences in protein copy numbers expressed from different genes in iPS cells, spanning over 7 orders of magnitude, from <100 to ∼100 million copies per cell (Fig. 1d). This wide dynamic range of expression means the 15 most abundant proteins represent >10% of the total protein molecules in the iPS cell (Fig.1e). These proteins include, as expected, histones, ribosomal proteins, translation factors and cytoskeletal components, which are abundant in most cell types. However, this hyper abundant group also includes two members of the glycolytic pathway, i.e. GAPDH (∼45 million copies) and ENO1 (∼18 million copies), along with the antioxidant PRDX1 (∼15 million copies), whose abundances are more variable in differentiated cell types.

The human iPSC proteome shows comprehensive coverage of known protein complexes, including subunits from 92% of all complexes described in the mammalian protein complex database CORUM^10^. It also includes multiple protein families involved in cell signalling. For example, we detect expression of 375 different protein kinases. This represents ∼74% of all confirmed human kinases^11^, consistent with predictions that hESCs express ∼300-400 kinases^12^. We also detect expression of 133 (∼70%) protein phosphatases^13^, 247 (∼66%) E3 ligases^14^ and 862 (∼53%) transcription factors^15^. Despite their functional importance, each of these protein families represent only a small portion of the total protein abundance in iPS cells: i.e. all kinases represent ∼1.17%, phosphatases ∼0.95%, E3 ligases ∼0.70% and transcription factors ∼ 2.28% of total protein copies.

### Proteomic profile of cell cycle, DNA repair and metabolism

#### Cell cycle

Human iPSCs are rapidly proliferating cells^16^. This is reflected in their proteome by high levels of key cell cycle regulators, including the D type cyclins, mitotic cyclins and DNA replication complexes. For example, all the MCM proteins (MCM2-7), which are the core components of the machinery that recognises DNA replication origins and licenses dormant origins^17^, are highly abundant (>1,3 million copies) in iPS cells. Conversely, the iPSC lines had low levels of the CDK inhibitors (CDKIs), which repress cell cycle progression (Fig. 2a). These observations help to explain how iPS cells achieve high rates of cell division.

**Figure 2.**
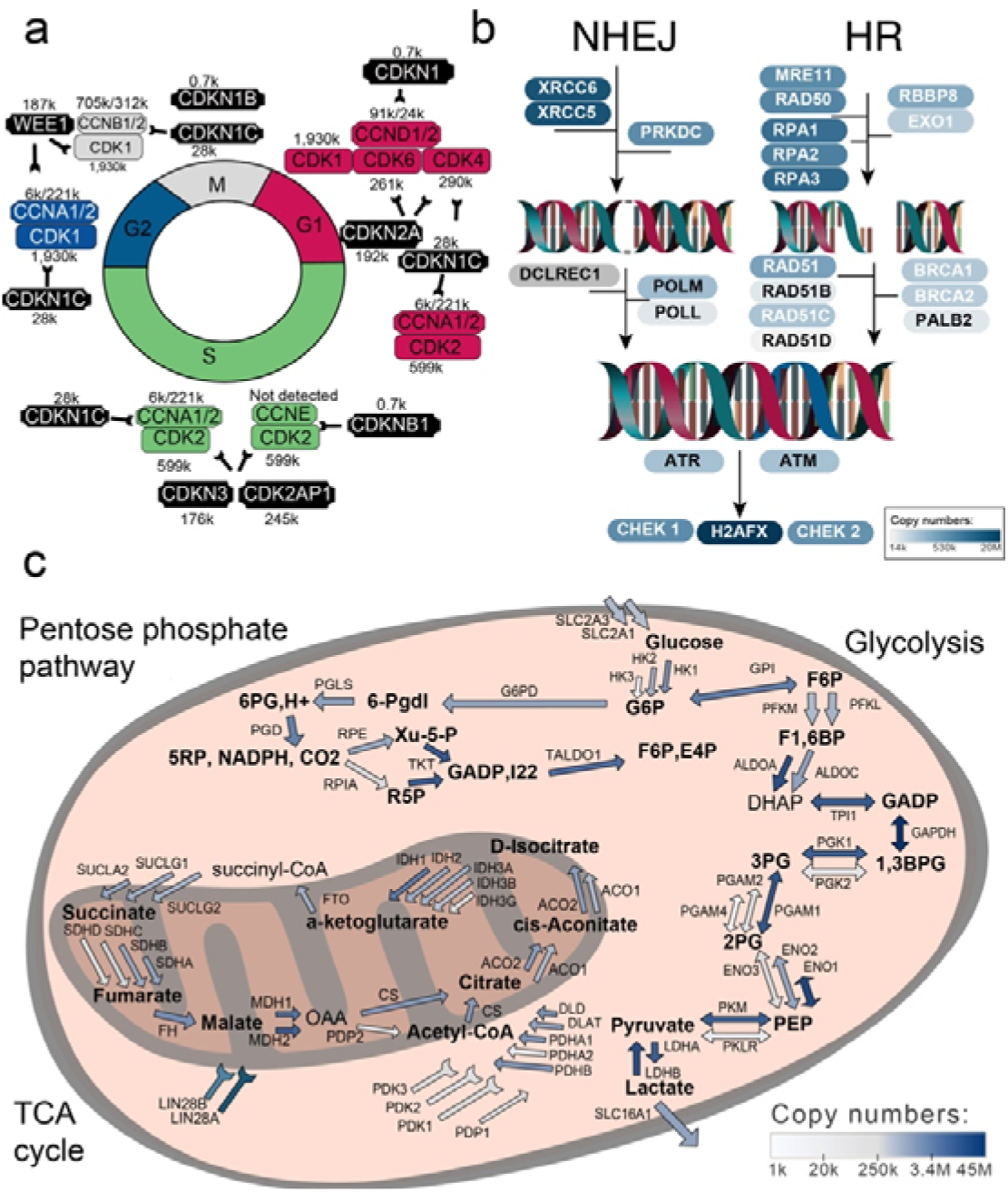
Cell cycle, DNA repair and metabolism: **(a)** Representation of the iPSC cell cycle, illustrating the phase duration, the phase specific cyclins, CDKs and their inhibitors. **(b)** iPSC metabolism, with the glycolytic, pentose phosphate and TCA pathways represented. Proteins are coloured in shades of blue to indicate copy numbers. **(c)** DNA double strand break pathways NEHJ and HR with proteins coloured in shades of blue to indicate copy numbers.

Increased expression of the CDK inhibitor CDKN1B is linked with lengthening of G1 and loss of pluripotency^18^. Consistent with the short G1 phase in iPSCs, our data show that the expression level of CDKN1B is low, (<1,000 copies per cell). CDKN1B was only detected in ∼3% of lines, compared with both cyclin D1 (∼91,000 copies per cell) and cyclin D2 (∼24,000 copies), which are detected in ∼70% and 95% of all lines, respectively. D Cyclins form a complex with CDK4/CDK6 and their inhibition lengthens G1 phase^19^.

Both CDK4 and CDK6 are significantly more abundant in iPSCs (>250,000 copies per cell), than their inhibitors, (e.g. CDKN1 & CDKN2). Thus, we detect two isoforms for CDKN2A expressed in iPSCs, p16INK4a and p12. The predominant isoform is p12, at ∼125,000 copies, which has been reported not to have activity towards CDK4/6^20^. The inhibitor CDKN1C is only present at ∼28,000 copies, i.e. ∼tenfold lower than CDK4 and CDK6. The CDK2 (∼599,000 copies)/CyclinA (∼227,000 copies) complex is also highly abundant. It contributes to a short G1 phase by phosphorylating and inactivating RB family proteins, which would otherwise bind to E2F4 and E2F5 to repress transcription of genes encoding proteins that regulate the G1-S phase transition^21^.

Moreover, CDK1 is the most abundant kinase within iPSCs with ∼2 million copies per cell. CDK1 plays a vital role in rapidly proliferating iPSCs as it drives both the G2-M and the G1-S transitions, via its association with cyclins D1, E and A^22^. The CDK1/CCNB1 complex also induces expression of LIN28A, which assists reprogramming to a glycolytic state by repressing oxidative phosphorylation (OXPHOS)^23^.

#### DNA Damage Repair

High rates of DNA replication and cell division are linked with an increased risk of DNA damage, which can lead to enhanced rates of mutation and/or cell death. A characteristic of cells responding to DNA damage is increased levels of induced DNA damage response factors. Interestingly, histone H2AFX, an indicator of DNA damage, arising from either Double Strand Breaks (DSB)^24^, or replicative stress^25^, is the 9th most abundant protein in iPSCs, with a median copy number of ∼20 million. However, it shows considerable variation in expression levels between the different iPSC lines, fluctuating between 16 and 74 million copies per cell.

A hallmark of pluripotent stem cells is their ability to differentiate into the three primary germ layers and thus any mutations they sustain could affect entire cell lineages during differentiation^26^. To avoid DNA damage being propagated to the progeny, stem cells have a robust DNA Damage Repair (DDR) system and show a lower mutation frequency than somatic cells. DSBs generated from reprogramming and replicative stress are mended by the intervention of the Homologous Recombination (HR) or Non-Homologous End Joining (NHEJ) pathways^27^. iPS cells are particularly effective at the error free repair of DSBs by HR, which is facilitated by a short G1 phase^28^. Proteins acting in both of these pathways are highly abundant in iPSCs (Fig. 2b). For example, RPA proteins are vital components of HR, they coat single-strand DNA to facilitate the loading of the RAD51 recombinase^24, 28^ and both RPA1 and 3 are present at >1.4 million copies per cell. Similarly, XRCCC6/5 (also known as Ku70 and Ku80), start the NHEJ process by forming a dimer on broken DNA ends and recruiting PRKDC (DNA-PK)^29^. PRKDC is the second most abundant kinase in iPS cells (∼1.2 million copies) and both XRCCC6/5 are present at >3 million copies per cell. Furthermore, all members of the DSB repair pathways are expressed at >14,000 copies per cell.

If DNA damage is not repaired, iPSCs can either temporarily arrest in G2 phase, upon ATM activation^30^, or activate cell death^31^. ATM will operate at DSBs and ATR at ssDNA stretches, phosphorylating the downstream targets CHEK1/2 and H2AFX. Efficient DDR is facilitated in iPS cells via high expression of SALL4, (∼500,000 copies), which favours ATM activation^32^ and DICER1, (∼145,000 copies), which helps resolve replicative stress^33^. CHEK1 and CHEK2 are among the top 10% most abundant kinases in iPS cells, (both>330,000 copies) and the p53 DNA damage induced transcription factor is also abundant, at ∼115,000 copies per cell. Vital Single Strand Breaks (SSB) repair proteins are also highly abundant with PARP1 with ∼3.7 million copies per cell and XRCC1 ∼350,000 copies.

In summary, we detect iPS cells expressing very high levels of a wide range of protein factors involved in the repair of DNA damage. This is consistent with the protection of iPSCs from an increased risk of DNA damage and mutation during rapid proliferation and subsequent differentiation.

#### Metabolism

The expression levels of key enzymes and molecules that control cell metabolism are highlighted in Fig. 2c, giving insights about the iPSC metabolic programs. All the iPSC lines expressed high levels of glycolytic enzymes. Glycolysis is inefficient for ATP production, compared with OXPHOS. Cells dependent upon glycolysis for energy metabolism therefore must sustain high levels of glucose uptake, which is reflected in the iPSC proteome by the high expression levels of multiple glucose transporters. For example, SLC2A1 (GLUT1) and SLC2A3 (GLUT3) are both present at ∼500,000 copies per cell in all the iPSC lines.

Within cells, glucose is converted to glucose 6-phosphate by Hexokinase, which is rate limiting for glucose metabolism. Cells with high glycolytic rates thus express elevated expression levels of Hexokinases (HKs)^34^. We detect abundant HK expression in iPSCs, with HK1 at ∼1,470,000 copies, followed by HK2 at ∼650,000 copies per cell. Multiple other glycolytic pathway components are also highly expressed in iPSCs (Fig. 2c), including GAPDH (∼45 million copies) and ENO1, PGAM1, PGK1 and PKM, each with >7 million copies.

Pyruvate produced by glycolysis can either be converted into L-lactate, via lactate dehydrogenases (LDH), or into Acetyl-CoA, via the pyruvate dehydrogenase complex (PDC)^35^. The lower reliance of iPSCs on OXPHOS, compared with somatic cells^36^, is consistent with the high expression of LDHs and the PDC inhibitors, PDK1-3 (Fig. 2c). The HipSci iPSC lines express high levels of both LDHA (∼7 million copies) and LDHB (∼13 million copies per cell). LDH isozymes have different affinity and inhibition patterns, with LDHB isozymes present in cells that are less inhibited by L-lactate^37^. Nonetheless L-lactate must still be removed by monocarboxylate transporters and our data show all iPSC lines have high levels (∼1 million copies) of the lactate transporter SLC16A1. We also detect high expression of specific amino acid transporters, such as LAT1 (SLC7A5) (∼635,000 copies) and its heavy subunit CD98 (SLC3A2) (∼1,500,000 copies).

We detect abundant expression in iPSCs of the key enzymes of the PDC and TCA cycle. The PDC, which converts Pyruvate to Acetyl-CoA and is linked with histone acetylation and maintenance of pluripotency, hence is still vital for iPSCs^38^. However, consistent with the lower reliance of iPSCs on OXPHOS, we also detect high expression of OXPHOS inhibitors, such as PDK1-3 and LIN28A/B^39^. Furthermore, we detect expression of all enzymes in the Pentose Phosphate Pathway (PPP). Congruent with reports that iPSCs have preference for the nonoxidative side of the PPP^40^, we detect highest levels of the non-oxidative enzymes Transketolase and Transaldolase, both expressed at >2 million copies per cell.

A feature of iPSCs is that they require high expression of antioxidants to reduce the oxidative stress caused by metabolism. In line with this requirement, our data show that the antioxidant PRDX1 is one of the most abundant proteins in all the iPSC lines, with ∼16 million copies per cell. PRDX1 has been shown to regulate gene expression in response to ROS^41^.

### Evaluating pluripotency and primed pluripotency markers

Human iPS cells share the hallmarks of primed pluripotency with their hESC counterparts, including the ability to self-renew and to differentiate into the three main germ layers. We initially evaluated the expression of 3 key factors known to maintain pluripotency and prevent spontaneous differentiation in primed iPSCs; i.e. SOX2, OCT4 and NANOG (Fig. 3a). All 3 were expressed ubiquitously across nearly all iPSC lines.

**Figure 3.**
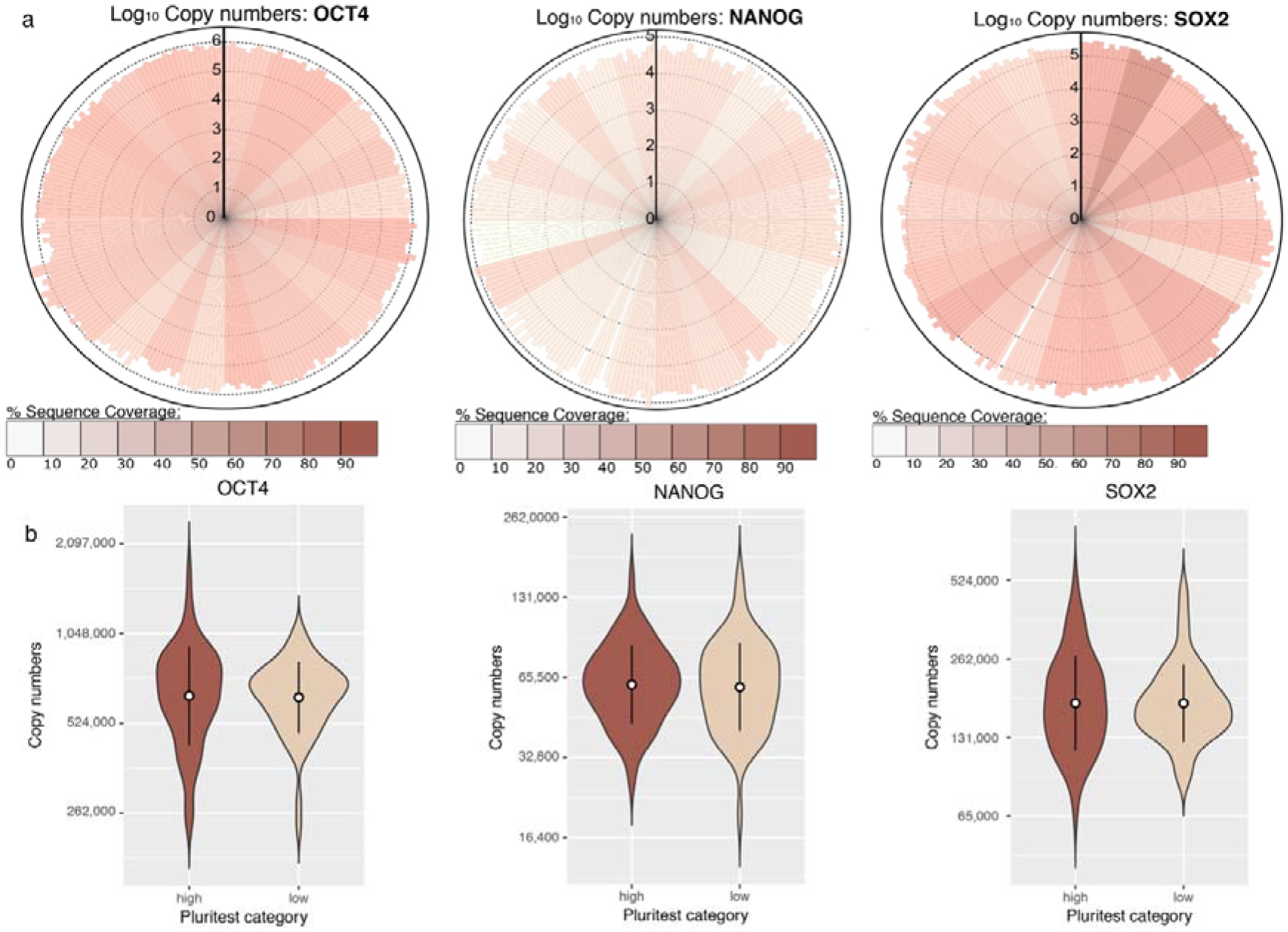
Evaluating iPSC pluripotency: **(a)** Radial bar plots showing Log_10_ copy numbers of SOX2, OCT4, and NANOG proteins in all HipSci lines. **(b)** Violin plots with geometric means and standard deviation showing the expression levels of SOX2, OCT4, and NANOG, when comparing ‘High’ vs ‘Low’ Pluritest score categories.

Focussing on iPSC lines derived from healthy donors, we used the Pluritest^3^ score to stratify the lines into ‘High’ and ‘Low’ Pluritest score categories (see methods). We compared protein expression across the ‘High’ and ‘Low’ populations for 123 lines. The levels of the 3 canonical pluripotency factors remained virtually identical between the two groups (Fig. 3b), consistent with the high QC within the HipSci pipeline^2^.

We evaluated proteomic changes within the two conditions and generated a Volcano plot comparing the ‘High’ vs ‘Low’ categories (Fig. 4a). The plot shows that the two populations are very similar. However, the statistical analysis revealed a subset of differentially expressed proteins affecting 3 main pathways, i.e. NF-kB, TGFB and RAS/RAF/ERK.

**Figure 4.**
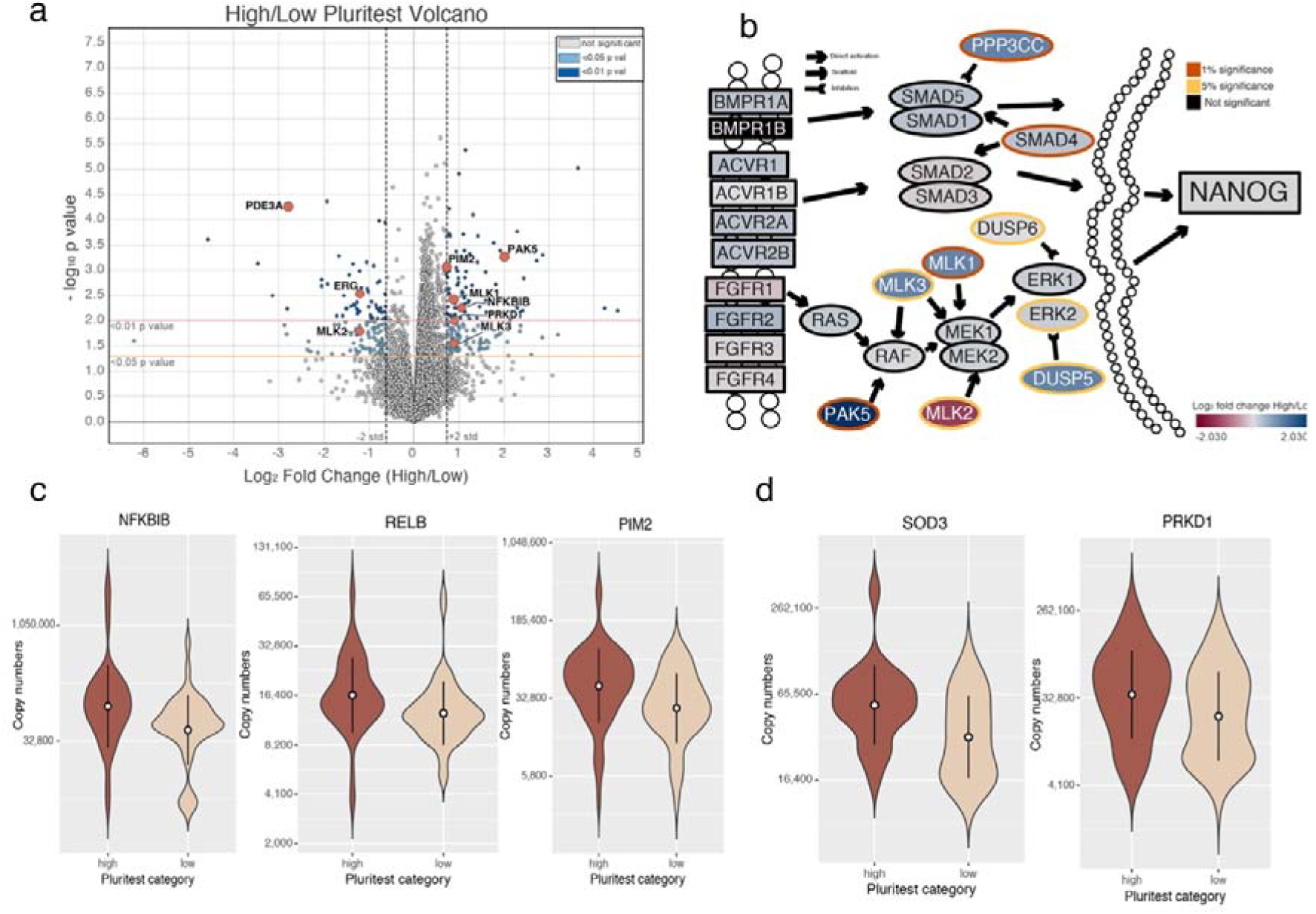
Fine-tuning primed pluripotency: **(a)** Volcano plot showing the Log_2_ fold change in protein abundance between High/Low Pluritest score vs the –log_10_ p-value for each protein detected in more than 2 TMT batches in both conditions and with more than 2 unique+razor peptides. **(b)** Activin/BMP/TGF-B and FGF2/ERK signalling pathways with components coloured by fold change and the borders by the p-value **(c)** Violin plots with geometric means and standard deviation showing the expression of NFKBIB, RELB and PIM2 across Pluritest categories. **(d)** Violin plots with geometric means and standard deviation showing the expression levels of the SOD3 and PRDK1 across Pluritest categories.

The canonical FGF2-RAS/RAF/ERK pathway is vital to primed pluripotency^42^ and our data show the core members remain virtually unchanged between populations. However, we detect increased expression of MLK1/3. MLKs can phosphorylate and activate MEK, with MLK1 and MLK3 being the most effective at activating ERK^43^. We also see an increase in the nuclear ERK phosphatase DUSP5, which is activated as a negative feedback control for ERK^44^ and PAK5, a constitutively active kinase that can phosphorylate BAD to prevent apoptosis^45^.

The Activin/TGFB pathway is important for primed pluripotency due to its direct modulation of NANOG expression^46^. Our data show that while canonical Activin/TGFB signalling remains unchanged between the two conditions (Fig. 4b), there is a difference with the non-canonical BMP4/TGF-B pathway. Though required to maintain pluripotency in mESC, BMP4 signalling promotes differentiation of primed PSCs^47^. Our data show increased expression in the ‘High’ Pluritest category of calcineurin, which is the catalytic subunit gamma of PP3 (Fig. 4b). FGF activated calcineurin can directly modulate BMP signalling by dephosphorylating SMAD1/SMAD5^48^.

Lastly, our data highlight multiple effects of antioxidants, including side-effects from the knockout serum replacement (KOSR) growth medium. KOSR is rich in vitamin C, which assists with the reprogramming of iPSCs^49^, however its effects extend beyond reprogramming. Vitamin C has been reported to have multiple effects on NF-kB signalling, including the inhibition of IKKβ and IKKα^50^. Our data show NFKBIB is not phosphorylated and degraded, suggesting reduced canonical NF-kB signalling. Another effect is seen through the alternative NF-kB signalling component, RELB/p52. In the ‘High’ Pluritest category we see an increase in RELB levels and its transcriptional target PIM2.

Furthermore, vitamin C promotes the expression SOD3, but not SOD1 or SOD2^51^. SOD3, which contributes to an antioxidative response by converting two superoxide radicals into hydrogen peroxide and water, is upregulated in cell lines in the ‘High’ Pluritest category. We also see upregulation of PRDK1, which is activated in response to ROS, more specifically by hydrogen peroxide^52^.

In summary, our data indicate that ‘High’ Pluritest category iPS cell lines have lower BMP4 signalling, a stronger antioxidant response and potentially less vulnerability to apoptotic signals.

### The Encyclopedia of Proteome Dynamics (EPD): Interactive iPSC analysis and visualisation

The EPD is an online database and web-application that provides open access to multi-omics data, featuring graphical navigation with interactive visualisations that enable data exploration in an intuitive, user-friendly manner^5^. All of the processed iPSC proteomics data, has been integrated within the EPD ecosystem and is available at: https://peptracker.com/epd/analytics/?section_id=40100.

As illustrated in Fig. 5, the EPD provides dynamic visualisations specifically created to analyse and explore the iPSC data. A dashboard presents an overview of the data for each specific protein, summarising correlation with RNA data, copy numbers and identification details (Fig. 5c). Global views of the dataset are provided with histograms (Fig. 5d) and bubble plots (Fig. 5e), both displaying the median protein copy numbers across all lines. For the Pluritest analysis, the Volcano plot (Fig. 5f), showing p-values and log2 fold changes, is available interactively. We also integrated the iPSC dataset with the Reactome^53^ Pathway widget for pathway analysis (Fig. 5g) and with the KinoViewer^8^, to explore kinase expression via the kinase phylogenetic tree (Fig. 4h).

**Figure 5.**
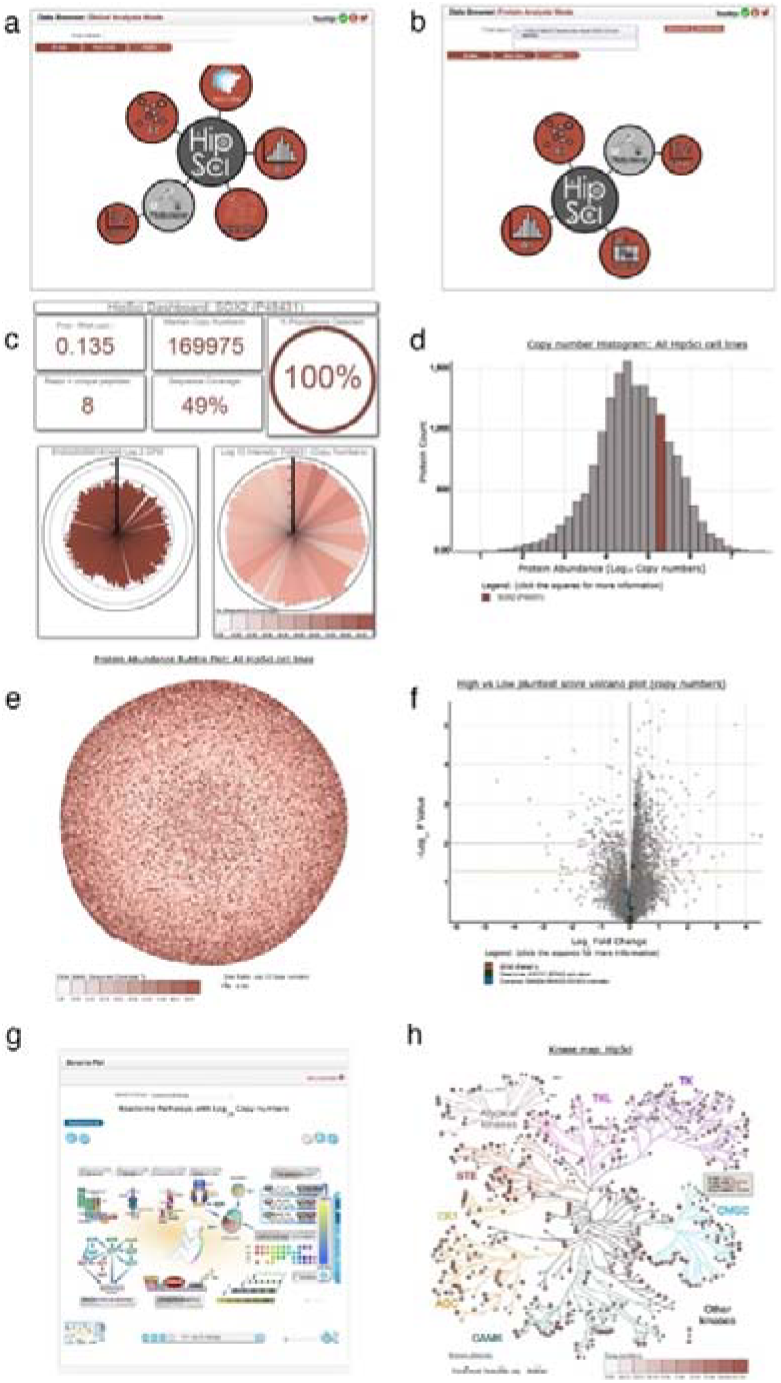
HipSci Data in the EPD: **(a)** Navigation interface for HipSci in the ‘Global Analysis Mode’. **(b)** Navigation interface for HipSci in the ‘Protein Analysis Mode’. **(c)** The HipSci dashboard. This can be accessed through the ‘Protein Analysis Mode’ and shows an overview of data available for the specific protein of interest. **(d)** Histogram of abundance for protein copy numbers across all iPS lines. A histogram can also be generated for each individual iPS line present within the HipSci dataset. **(e**) Bubble plot of protein abundance calculated for all iPS cell lines, with sequence coverage shown on the colour scale. It can be generated for either raw copy numbers, or Log_10_ transformed copy numbers. It can also be generated separately for each iPS line present within the HipSci dataset. **(f)** Volcano plot showing the comparison of protein expression in iPS lines with ‘High’ vs ‘Low’ Pluritest scores. Highlighted elements display the protein, Pathway and Protein Complex search functionality. **(g)** Reactome pathway analysis showing ‘Developmental Biology’ pathway, with HipSci copy numbers overlaid. **(h)** Interactive Kinase map showing kinase expression in iPS cells with copy numbers overlaid via the colour scale.

### hiPSC spectral library

To facilitate further MS analyses on human iPS cells, we also created a data independent acquisition (DIA) Spectral Library. To generate the library, an initial DDA workflow was set up (Fig. 6). Three representative iPSC lines were selected for in-depth analysis, via reversed-phase and HILIC chromatography. 24 fractions were analysed by LC-MS/MS on a Q-Exactive Plus Orbitrap mass-spectrometer in ‘Label Free’ mode, with two technical replicates per line. The samples were spiked with the Biognosys IRT Kit to align retention time. In total, 288 raw files were collected and quantified using MaxQuant v 1.6.0.13. This output was used to generate the spectral library using Spectronaut.

**Figure 6.**
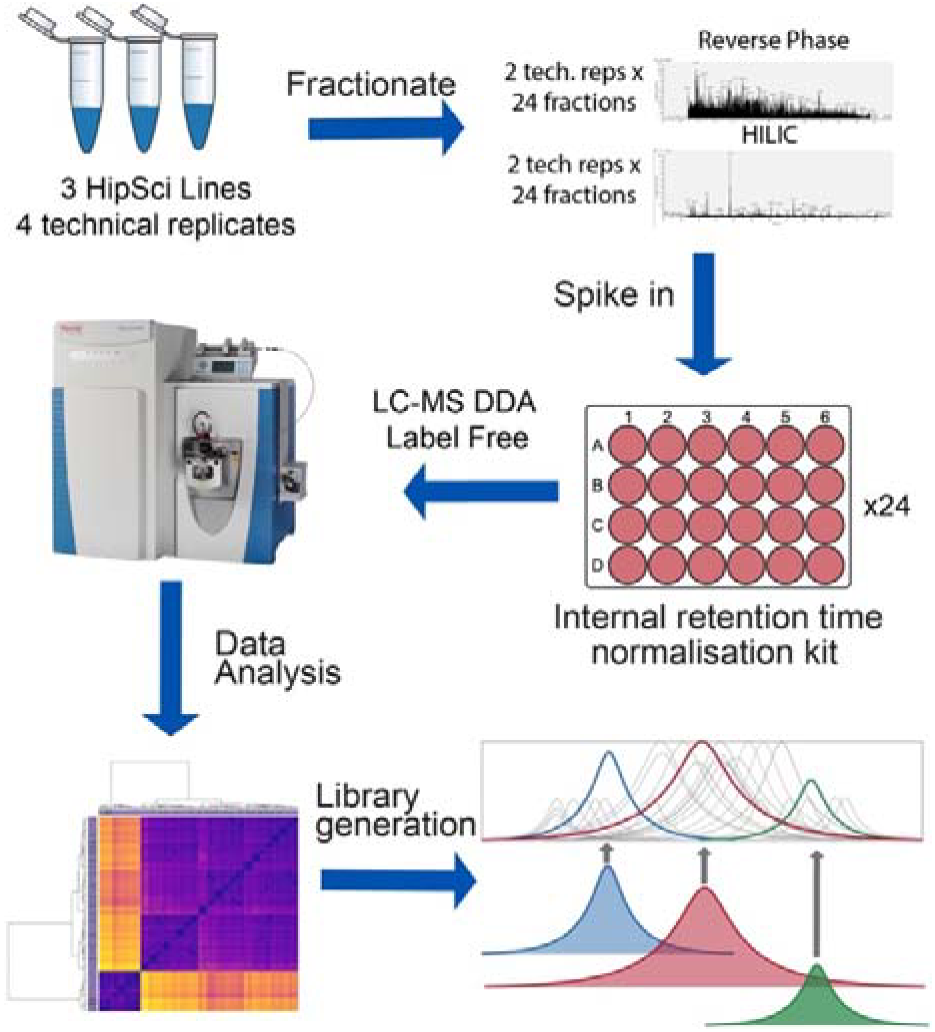
DIA Spectral Library: Workflow showing the process used to generate the DIA library specific for human iPSCs.

The resulting human iPSC DIA library is compatible with Orbitrap MS platforms and with popular DIA software packages, e.g. Skyline and Spectronaut. This iPSC DIA library is freely available within the PRIDE submission (PXD010557).

## Discussion

Human induced pluripotent stem cells have great potential for therapeutic and research applications, providing improved ethical models for studies on disease mechanisms and personalised regenerative medicine. We provide here a valuable research resource with the most in-depth proteome map of human iPSCs reported to date.

Our proteomic data provides insights into iPSC metabolism, highlighting the important role of glycolysis, along with elevated levels of glucose, amino acid and lactate transporters. The hyper-abundant group of iPSC proteins include the glycolytic pathway members GAPDH and ENO1, along with the elongation factor EEF1A1 and the antioxidant PRDX1. Their high expression highlights mechanisms supporting rapid proliferation and high metabolic activity in iPS cells. The proteomic profile of metabolic enzymes in iPS cells revealed high expression of OXPHOS inhibitors, e.g. PDK1-3 and LIN28A/B, as well as showing a preference for the anaerobic side of the PPP.

Human stem cells are characterised by a short G1 phase, with >50% of asynchronous hESCs in S phase^54^. Lengthening of G1 within pluripotent stem cells (PSCs) is linked with a loss of pluripotency^55^. Our data revealed high expression levels of G1 modulating CDKs (with CDK1 being the most abundant kinase) and their respective cyclins, along with correspondingly low expression levels of their respective inhibitor proteins. This accounts for the observed short G1 phase in iPS cells.

The proteomic data also help to explain the highly efficient DNA Damage Repair phenotype of iPS cells, which is important to prevent accumulation of mutations during proliferation and differentiation. Thus, we detect high expression levels of members of the NHEJ and HR pathways, along with elevated expression of p53, which will trigger cell death should DNA damage not be corrected.

The HipSci pipeline included in the QC process an evaluation also of the pluripotent potential of each line, reflected in a ‘Pluritest Score’^2^. We used these Pluritest scores measured for the individual iPSC lines to correlate proteome variation with pluripotency. This identified new potential protein regulators of the primed pluripotent state. For example, in lines with high Pluritest scores, we detected elevated levels of PPP3CC, which modulates BMP4 signalling. We also detected high levels of PAK5, which has been shown to prevent apoptosis, both by phosphorylating BAD and by activating and translocating RAF1 to mitochondria^56^. Overall, our data support the view that the ‘High’ Pluritest category is linked potentially with reduced BMP4 signalling and increased resistance to apoptotic signals. Moreover, the data revealed an antioxidant response to the growth medium, with reduced canonical NF-kB signalling and increased expression of the antioxidant proteins SOD3 and PRDK1.

To add value to these rich proteomic data, we have integrated all of the protein-level information into the Encyclopedia of Proteome Dynamics, an open access, interactive online web app. Additionally, all of the corresponding raw MS data are provided for open access via the ProteomeXchange PRIDE repository (PXD010557). Furthermore, we have generated a human iPSC spectral library, to support future proteomic studies on healthy and disease iPSCs lines using data independent acquisition (DIA). The iPSC spectral library is also freely available via the PRIDE repository. We are confident these data will provide a valuable resource that will facilitate further research and applications using human iPSCs.

## Methods

### Generation of iPSC lines

All lines included in this study are part of the HipSci resource and were reprogrammed from primary fibroblasts as previously described^2^. Out of the total of more than 800 iPSC lines available within the HipSci resource (www.hipsci.org), 217 lines, predominantly from healthy donors, were selected for in depth proteomic analysis in this study using Tandem Mass Tag Mass Spectrometry.

### TMT Sample preparation

For protein extraction, iPSC cell pellets were washed with ice cold PBS and redissolved immediately in 200 μL of lysis buffer (8 M urea in 100 mM triethyl ammonium bicarbonate (TEAB)) and mixed at room temperature for 15 minutes. The DNA content of the cells is sheared using ultrasonication (6 X 20 s on ice). The proteins were reduced using tris-carboxyethylphosphine TCEP (25 mM) for 30 minutes at room temperature, then alkylated in the dark for 30 minutes using iodoacetamide (50 mM). Total protein was quantified using the EZQ assay (Life Technologies). The lysates were diluted with 100 mM TEAB 4-fold for the first digestion with mass spectrometry grade lysyl endopeptidase, Lys-C (Wako, Japan), then further diluted 2.5-fold before a second digestion with trypsin. Lys-C and trypsin were used at an enzyme to substrate ratio of 1:50 (w/w). The digestions were carried out overnight at 37ºC, then stopped by acidification with trifluoroacetic acid (TFA) to a final concentration of 1% (v:v). Peptides were desalted using C18 Sep-Pak cartridges (Waters) following manufacturer’s instructions.

For tandem mass tag (TMT)-based quantification, the dried peptides were re-dissolved in 100 mM TEAB (50 μl) and their concentration was measured using a fluorescent assay (CBQCA, Life Technologies). 100 μg of peptides from each cell line to be compared, in 100 μl of TEAB, were labelled with a different TMT tag (20 μg ml^−1^ in 40 μl acetonitrile) (Thermo Scientific), for 2 h at room temperature. After incubation, the labelling reaction was quenched using 8 μl of 5% hydroxylamine (Pierce) for 30 min and the different cell lines/tags were mixed and dried *in vacuo*.

The TMT samples were fractionated using off-line high-pH reverse-phase (RP) chromatography: samples were loaded onto a 4.6 × 250 mm Xbridge BEH130 C18 column with 3.5-μm particles (Waters). Using a Dionex bioRS system, the samples were separated using a 25-min multistep gradient of solvents A (10 mM formate at pH 9) and B (10 mM ammonium formate pH 9 in 80% acetonitrile), at a flow rate of 1 ml min^−1^. Peptides were separated into 48 fractions, which were consolidated into 24 fractions. The fractions were subsequently dried and the peptides re-dissolved in 5% formic acid and analysed by LC–MS/MS.

### TMT LC–MS/MS

*TMT-based analysis.* Samples were analysed using an Orbitrap Fusion Tribrid mass spectrometer (Thermo Scientific), equipped with a Dionex ultra-high-pressure liquid-chromatography system (RSLCnano). RPLC was performed using a Dionex RSLCnano HPLC (Thermo Scientific). Peptides were injected onto a 75 µm × 2 cm PepMap-C18 pre-column and resolved on a 75 µm × 50 cm RP-C18 EASY-Spray temperature-controlled integrated column-emitter (Thermo Scientific), using a four-hour multistep gradient from 5% B to 35% B with a constant flow of 200 nl min^−1^. The mobile phases were: 2% ACN incorporating 0.1% FA (solvent A) and 80% ACN incorporating 0.1% FA (solvent B). The spray was initiated by applying 2.5 kV to the EASY-Spray emitter and the data were acquired under the control of Xcalibur software in a data-dependent mode using top speed and 4 s duration per cycle. The survey scan is acquired in the orbitrap covering the *m*/*z* range from 400 to 1,400 Thomson with a mass resolution of 120,000 and an automatic gain control (AGC) target of 2.0 x 10^5^ ions. The most intense ions were selected for fragmentation using CID in the ion trap with 30% CID collision energy and an isolation window of 1.6 Th. The AGC target was set to 1.0 x 10^4^ with a maximum injection time of 70 ms and a dynamic exclusion of 80 s.

During the MS3 analysis for more accurate TMT quantifications, 5 fragment ions were co-isolated using synchronous precursor selection using a window of 2 Th and further fragmented using HCD collision energy of 55%. The fragments were then analysed in the orbitrap with a resolution of 60,000. The AGC target was set to 1.0 x 10^5^ and the maximum injection time was set to 105 ms.

### Accession codes

All of the mass-spectrometry data generated from the TMT batches, the DIA library, fasta file, and MaxQuant outputs have been uploaded to PRIDE (PXD10557).

### Identification & Quantification

The TMT-labelled samples were collected and analysed using Maxquant^7, 57^ v. 1.6.0.13 in a single large run. The FDR threshold was set to 5% on the Peptide Spectrum Match, Peptide and Protein level. Proteins and peptides were identified using the UniProt human reference proteome database (SwissProt & TrEMBL). Run parameters have been deposited to PRIDE^5^ along with the full MaxQuant quantification output (PDX010557).

Data for the analysis were obtained from the ProteinGroups.txt output of MaxQuant. Contaminants, reverse hits and ‘only identified by site’ were excluded from analysis. For either a protein, or a protein group to be considered, we required at least one unique peptide mapping to it. Overall, we quantified 16,755 protein groups in at least one of the samples.

### Copy number generation

Every TMT batch had 1 control cell line (bubh3, reporter channel 0) and 9 sample cell lines. Protein copy numbers were calculated following the proteomic ruler^9^ and using the MS3 intensity. To minimise potential batch effects, the protein copy numbers were normalised using the reference cell line present in every batch.

For every protein, the references lines in each TMT batch were used to calculate a median copy number, and then calculate a ratio between the control in every TMT batch and the median. All values within the batch were corrected via this ratio.

### Lead razor protein assignment

The lead razor protein for each Protein Group was modified from the MaxQuant output. The number of peptides that could theoretically be assigned to each element of the protein groups were selected, and all the elements of the Protein Group with highest number of peptides were selected. If there was only one candidate protein, then this element became the lead razor protein, if multiple candidates were present additional filtering was required.

Among the candidates the priority was given to the canonical SwissProt entry, if no canonical entry was present it would default to a reviewed protein isoform. If no reviewed proteins isoforms were present among the candidate proteins, then the lead element would be assigned to a TrEMBL proteoform. A TrEMBL proteoform would only be assigned a lead protein role based on the previously described scenario.

### Chromosome mapping

To map gene products to their specific chromosomes, we utilised the UniProt^58^ protein-chromosome mapping file. We used the file to produce a list of unique protein coding genes for each specific chromosome. Subsequently, we mapped the proteins detected in our iPSC dataset to their corresponding chromosomes based on the UniProt mapping file and produced a list of genes for each chromosome as well. We compared the iPSC specific list of genes, with the reference list to determine the percentage of protein coding genes detected in our iPSC dataset for each chromosome.

### Statistical analysis

The Pluritest statistical analysis, illustrated on the volcano plot, was generated based on the copy number and Pluritest scores data. The lines were stratified into two categories, i.e. ‘High’ and ‘Low’, based on the median Pluritest score. Only proteins that were detected in at least two distinct TMT batches for each condition were selected for analysis. Additionally, 2 or more Unique and Razor peptides had to be assigned to the protein group.

P-values were calculated in R utilising the bioconductor package Linear Models for Microarray Data (LIMMA) version 3.7. The fold change used was also calculated by LIMMA and uses the Log_2_ geometric mean. Q-values were generated in R using the “qvalue” package version 2.10.0

### Dynamic visualisations within the EPD

All processed proteomic data were integrated into the Encyclopedia of Proteome Dynamics^6^ (https://peptracker.com/epd/analytics/). The tabular data are stored within Cassandra, and the relationships between datasets and identified proteins were modelled and stored within Neo4j.

The EPD server runs on Django, while the front end is a mixture of Angular, jQuery and D3.js. All of the visualisations were generated via D3.js as a client-side JavaScript library. The Reactome^53^ Pathway browser was integrated into the EPD using the Reactome JavaScript widget and Analysis Services.

## Acknowledgements

We thank all of our collaborators who worked on the HipSci project. We would also like to thank our University of Dundee colleagues, Greg Findlay, Anton Gartner and Doreen Cantrell, for helpful discussions relevant to iPS cell signalling and metabolism.

## Author contributions

A.I.L. D.B. and A.B. conceived the study. D.B & A.I.L. supervised the project. V.A. assisted with the sample tracking and metadata storage. J.H. assisted in the interpretation of the data. D.B. designed and performed all proteomics experiments and created the DIA library. A.B. integrated the data into the EPD and created the visualisations. A.B. performed the data analysis and interpretation. The paper was written by A.B. and D.B. and edited by all authors.

## References

1. Takahashi, K. et al. Induction of pluripotent stem cells from adult human fibroblasts by defined factors. Cell 131, 861–872 (2007).

2. Kilpinen, H. et al. Common genetic variation drives molecular heterogeneity in human iPSCs. Nature 546, 370–375 (2017).

3. Muller, F.J. et al. A bioinformatic assay for pluripotency in human cells. Nat Methods 8, 315–317 (2011).

4. Vizcaino, J.A. et al. ProteomeXchange provides globally coordinated proteomics data submission and dissemination. Nat Biotechnol 32, 223–226 (2014).

5. Vizcaino, J.A. et al. 2016 update of the PRIDE database and its related tools. Nucleic Acids Res 44, D447–456 (2016).

6. Brenes, A., Afzal, V., Kent, R. & Lamond, A.I. The Encyclopedia of Proteome Dynamics: a big data ecosystem for (prote)omics. Nucleic Acids Res 46, D1202–D1209 (2018).

7. Cox, J. & Mann, M. MaxQuant enables high peptide identification rates, individualized p.p.b.-range mass accuracies and proteome-wide protein quantification. Nat Biotechnol 26, 1367–1372 (2008).

8. Zubarev, R.A. The challenge of the proteome dynamic range and its implications for in-depth proteomics. Proteomics 13, 723–726 (2013).

9. Wisniewski, J.R., Hein, M.Y., Cox, J. & Mann, M. A “proteomic ruler” for protein copy number and concentration estimation without spike-in standards. Mol Cell Proteomics 13, 3497–3506 (2014).

10. Ruepp, A. et al. CORUM: the comprehensive resource of mammalian protein complexes--2009. Nucleic Acids Res 38, D497–501 (2010).

11. Brenes, A. & Lamond, A.I. The Encyclopedia of Proteome Dynamics: The KinoViewer. Bioinformatics (2018).

12. Fernandez-Alonso, R., Bustos, F., Williams, C.A.C. & Findlay, G.M. Protein Kinases in Pluripotency-Beyond the Usual Suspects. J Mol Biol 429, 1504–1520 (2017).

13. Chen, M.J., Dixon, J.E. & Manning, G. Genomics and evolution of protein phosphatases. Sci Signal 10 (2017).

14. Medvar, B., Raghuram, V., Pisitkun, T., Sarkar, A. & Knepper, M.A. Comprehensive database of human E3 ubiquitin ligases: application to aquaporin-2 regulation. Physiol Genomics 48, 502–512 (2016).

15. Lambert, S.A. et al. The Human Transcription Factors. Cell 172, 650–665 (2018).

16. Pauklin, S. & Vallier, L. The cell-cycle state of stem cells determines cell fate propensity. Cell 155, 135–147 (2013).

17. Blow, J.J., Ge, X.Q. & Jackson, D.A. How dormant origins promote complete genome replication. Trends Biochem Sci 36, 405–414 (2011).

18. Menchon, C., Edel, M.J. & Izpisua Belmonte, J.C. The cell cycle inhibitor p27Kip(1) controls self-renewal and pluripotency of human embryonic stem cells by regulating the cell cycle, Brachyury and Twist. Cell Cycle 10, 1435–1447 (2011).

19. Dong, P., Zhang, C., Parker, B.T., You, L. & Mathey-Prevot, B. Cyclin D/CDK4/6 activity controls G1 length in mammalian cells. PLoS One 13, e0185637 (2018).

20. Poi, M.J. et al. Evidence that P12, a specific variant of P16(INK4A), plays a suppressive role in human pancreatic carcinogenesis. Biochem Biophys Res Commun 436, 217–222 (2013).

21. Conklin, J.F., Baker, J. & Sage, J. The RB family is required for the self-renewal and survival of human embryonic stem cells. Nat Commun 3, 1244 (2012).

22. Hu, X. & Moscinski, L.C. Cdc2: a monopotent or pluripotent CDK? Cell Prolif 44, 205–211 (2011).

23. Wang, X.Q. et al. CDK1-PDK1-PI3K/Akt signaling pathway regulates embryonic and induced pluripotency. Cell Death Differ 24, 38–48 (2017).

24. Turinetto, V. et al. High basal gammaH2AX levels sustain self-renewal of mouse embryonic and induced pluripotent stem cells. Stem Cells 30, 1414–1423 (2012).

25. Zeman, M.K. & Cimprich, K.A. Causes and consequences of replication stress. Nat Cell Biol 16, 2–9 (2014).

26. Behrens, A., van Deursen, J.M., Rudolph, K.L. & Schumacher, B. Impact of genomic damage and ageing on stem cell function. Nat Cell Biol 16, 201–207 (2014).

27. Tilgner, K. et al. A human iPSC model of Ligase IV deficiency reveals an important role for NHEJ-mediated-DSB repair in the survival and genomic stability of induced pluripotent stem cells and emerging haematopoietic progenitors. Cell Death Differ 20, 1089–1100 (2013).

28. Vitale, I., Manic, G., De Maria, R., Kroemer, G. & Galluzzi, L. DNA Damage in Stem Cells. Mol Cell 66, 306–319 (2017).

29. Ma, C.J., Gibb, B., Kwon, Y., Sung, P. & Greene, E.C. Protein dynamics of human RPA and RAD51 on ssDNA during assembly and disassembly of the RAD51 filament. Nucleic Acids Res 45, 749–761 (2017).

30. Stambrook, P.J. & Tichy, E.D. Preservation of genomic integrity in mouse embryonic stem cells. Adv Exp Med Biol 695, 59–75 (2010).

31. Marechal, A. et al. PRP19 transforms into a sensor of RPA-ssDNA after DNA damage and drives ATR activation via a ubiquitin-mediated circuitry. Mol Cell 53, 235–246 (2014).

32. Xiong, J. et al. Stemness factor Sall4 is required for DNA damage response in embryonic stem cells. J Cell Biol 208, 513–520 (2015).

33. Swahari, V. et al. Essential Function of Dicer in Resolving DNA Damage in the Rapidly Dividing Cells of the Developing and Malignant Cerebellum. Cell Rep 14, 216–224 (2016).

34. Bustamante, E., Morris, H.P. & Pedersen, P.L. Energy metabolism of tumor cells. Requirement for a form of hexokinase with a propensity for mitochondrial binding. J Biol Chem 256, 8699–8704 (1981).

35. Teslaa, T. & Teitell, M.A. Pluripotent stem cell energy metabolism: an update. EMBO J 34, 138–153 (2015).

36. Wu, J., Ocampo, A. & Belmonte, J.C.I. Cellular Metabolism and Induced Pluripotency. Cell 166, 1371–1385 (2016).

37. Rogatzki, M.J., Ferguson, B.S., Goodwin, M.L. & Gladden, L.B. Lactate is always the end product of glycolysis. Front Neurosci 9, 22 (2015).

38. Moussaieff, A. et al. Glycolysis-mediated changes in acetyl-CoA and histone acetylation control the early differentiation of embryonic stem cells. Cell Metab 21, 392–402 (2015).

39. Zhang, J. et al. LIN28 Regulates Stem Cell Metabolism and Conversion to Primed Pluripotency. Cell Stem Cell 19, 66–80 (2016).

40. Varum, S. et al. Energy metabolism in human pluripotent stem cells and their differentiated counterparts. PLoS One 6, e20914 (2011).

41. Hopkins, B.L. et al. A Peroxidase Peroxiredoxin 1-Specific Redox Regulation of the Novel FOXO3 microRNA Target let-7. Antioxid Redox Signal 28, 62–77 (2018).

42. Lotz, S. et al. Sustained levels of FGF2 maintain undifferentiated stem cell cultures with biweekly feeding. PLoS One 8, e56289 (2013).

43. Marusiak, A.A. et al. Mixed lineage kinases activate MEK independently of RAF to mediate resistance to RAF inhibitors. Nat Commun 5, 3901 (2014).

44. Buffet, C. et al. DUSP5 and DUSP6, two ERK specific phosphatases, are markers of a higher MAPK signaling activation in BRAF mutated thyroid cancers. PLoS One 12, e0184861 (2017).

45. Zhong, J., Troppmair, J. & Rapp, U.R. Independent control of cell survival by Raf-1 and Bcl-2 at the mitochondria. Oncogene 20, 4807–4816 (2001).

46. Xu, R.H. et al. NANOG is a direct target of TGFbeta/activin-mediated SMAD signaling in human ESCs. Cell Stem Cell 3, 196–206 (2008).

47. Feng, L. et al. Discovery of a Small-Molecule BMP Sensitizer for Human Embryonic Stem Cell Differentiation. Cell Rep 15, 2063–2075 (2016).

48. Cho, A. et al. Calcineurin signaling regulates neural induction through antagonizing the BMP pathway. Neuron 82, 109–124 (2014).

49. Esteban, M.A. & Pei, D. Vitamin C improves the quality of somatic cell reprogramming. Nat Genet 44, 366–367 (2012).

50. Carcamo, J.M. et al. Vitamin C is a kinase inhibitor: dehydroascorbic acid inhibits IkappaBalpha kinase beta. Mol Cell Biol 24, 6645–6652 (2004).

51. Singh, B. & Bhat, H.K. Superoxide dismutase 3 is induced by antioxidants, inhibits oxidative DNA damage and is associated with inhibition of estrogen-induced breast cancer. Carcinogenesis 33, 2601–2610 (2012).

52. Waldron, R.T. & Rozengurt, E. Oxidative stress induces protein kinase D activation in intact cells. Involvement of Src and dependence on protein kinase C. J Biol Chem 275, 17114–17121 (2000).

53. Fabregat, A. et al. The Reactome pathway Knowledgebase. Nucleic Acids Res 44, D481–487 (2016).

54. Hindley, C. & Philpott, A. The cell cycle and pluripotency. Biochem J 451, 135–143 (2013).

55. Zhu, H., Hu, S. & Baker, J. JMJD5 regulates cell cycle and pluripotency in human embryonic stem cells. Stem Cells 32, 2098–2110 (2014).

56. Wu, X., Carr, H.S., Dan, I., Ruvolo, P.P. & Frost, J.A. p21 activated kinase 5 activates Raf-1 and targets it to mitochondria. J Cell Biochem 105, 167–175 (2008).

57. Tyanova, S., Temu, T. & Cox, J. The MaxQuant computational platform for mass spectrometry-based shotgun proteomics. Nat Protoc 11, 2301–2319 (2016).

58. The UniProt, C. UniProt: the universal protein knowledgebase. Nucleic Acids Res 45, D158–D169 (2017).

